# Integrated Gait and Pose Analysis Utilizing Computer Vision for Parkinsonian Behavioral Phenotyping in Mice

**DOI:** 10.64898/2026.03.10.706230

**Authors:** Matthew J. Jennings, Audrey Anigbo, Serge Przedborski

## Abstract

Synucleinopathies can be biologically advanced before overt parkinsonism is clinically apparent, highlighting the need for objective, sensitive motor endpoints. We examined the mThy1-α-synuclein line 61 (L61-Tg) mouse, which shows progressive synucleinopathy with early circuit dysfunction, using an integrated pipeline combining CatWalk XT gait analysis and markerless pose estimation from the same CatWalk videos. Two cohorts of male L61-Tg and nontransgenic littermates were assessed at 12 and 18 months. DeepLabCut tracking of four landmarks showed highest accuracy at the tail base. We thus quantified mediolateral instability as within-run variance of tail-base lateral position. L61-Tg mice exhibited increased tail-base lateral variance at both ages. CatWalk mixed-effects modeling identified six genotype-dependent parameters at 12 months, and a progressive increase in hind base of support at 18 months. Comparison across measures showed that discrimination between L61-Tg and non-transgenic was similarly high for hind base of support and tail-base lateral instability the two were nonetheless synergistic, and the approaches are therefore complementary to one-another in the determination of synucleinopathy motor phenotypes. This combined gait-pose strategy provides scalable, interpretable endpoints for preclinical Parkinson-like phenotyping and therapeutic testing.

## INTRODUCTION

Synucleinopathies, including Parkinson’s disease (PD), Parkinson’s disease dementia (PDD), and dementia with Lewy bodies (DLB), are unified by the misfolding and accumulation of α-synuclein and the gradual failure of neural circuits that sustain normal movement and cognition^1^. Increasingly, prodromal features such as hyposmia and REM sleep behavior disorder (RBD), especially when paired with emerging α-synuclein-based biomarkers, can support the presence of an underlying synucleinopathy well before a patient meets clinical criteria for parkinsonism^2^. Yet what remains underdeveloped are objective, quantitative, early motor readouts that (i) sensitively capture the transition from prodrome to overt parkinsonism (“phenoconversion”) and can be used as practical endpoints to monitor response to experimental disease-modifying therapies. Critically, this prodromal window is where disease modification is most plausible, but where our outcome measures are often least informative. For a neuropathologic framing of synucleinopathies, see Koga et al^1^ for the clinical spectrum of Lewy body dementias including PDD and DLB, see Walker et al^3^.

To interrogate this brain-behavior dissociation, we focus on the mThy1-α-synuclein “line 61” transgenic mouse (hereafter termed L61-Tg), which overexpresses human wild-type α-synuclein in neurons under the Thy1 promoter^4^. This genetic strategy aims at modeling human synucleinopathy biology, where increased SNCA dosage (e.g., locus duplication) can cause autosomal-dominant parkinsonism often complicated by dementia^5^. In L61-Tg mice, α-synuclein pathology develops broadly across vulnerable circuits and is accompanied by progressive perturbations in nigrostriatal dopamine physiology, changes in extracellular dopamine tone and synaptic modulation that precede later declines in striatal dopamine content^6^. Importantly, this model also expresses a gradual, age-dependent behavioral syndrome with early sensorimotor anomalies on challenging tasks^7^, while more overt deficits in conventional motor readouts emerge later, replicating a key feature of synucleinopathies wherein pathology and circuit dysfunction occur long before “classic” motor disability is obvious.

However, despite widespread use of L61-Tg mice, the published description of gait and its evolution remain limited. Challenging assays (e.g., beam-based paradigms) can reveal early performance deficits^8^, but such tasks blur the line between primary gait impairment and contributions from balance, motivation, sensory integration, or executive control, domains that are also affected across the PD-PDD-DLB spectrum. What is required are sensitive, scalable approaches that can quantify gait *as gait*, and do so in a way that is both data-rich and resistant to observer bias.

Automated gait analysis platforms address part of this need. The CatWalk XT system provides high-throughput spatiotemporal and interlimb coordination metrics and has been validated in toxin-based parkinsonian mouse models (e.g., MPTP) where CatWalk parameters track dopaminergic injury severity^9^. Moreover, computational methods can extract still more signal from CatWalk outputs: a machine learning pipeline has demonstrated that gait features can discriminate biologically meaningful experimental groups^10^, underscoring how much information is embedded in step sequences when we analyze them at scale.

However, CatWalk alone is not a complete description of movement. It captures footprints and derived gait parameters, but it does not natively resolve whole-body pose, information that can be critical for interpreting gait in synucleinopathy models (for example, whether a change in stride reflects bradykinesia, postural instability, compensatory trunk strategy, or altered tail/head dynamics). Here, markerless pose estimation offers complementarity. DeepLabCut (DLC) enables accurate tracking of user-defined body parts without physical markers^11^, reducing experimenter burden while allowing animals to move more naturally. Markerless approaches are not merely convenient, they help avoid experimental artifacts associated with marker placement and handling; indeed, in rodent locomotion studies, marker-based methods can introduce practical and behavioral confounds that motivate markerless alternatives (e.g., marker-related alteration of natural gait is discussed in the context of markerless kinematics^12^).

Here, we asked whether simultaneous CatWalk-based gait analysis and DLC-based markerless pose tracking can detect and characterize motor deficits in L61-Tg mice earlier than conventional behavioral readouts, and whether this richer phenotype clarifies how α-synuclein overexpression shapes the relationship between gait, posture, and progressive nigrostriatal dysfunction.

We therefore combine CatWalk XT with DLC to create a unified gait-pose phenotype for the L61-Tg model. CatWalk provides standardized, high-dimensional gait metrics; DLC contributes complementary head/tail (and potentially trunk) kinematics that anchor those gait metrics to posture and body strategy. This integrated, automated pipeline is designed to increase sensitivity to motor abnormalities that may precede overt impairment, improve reproducibility by minimizing observer-dependent scoring, and yield interpretable, quantitative endpoints suited for therapeutic testing and genetic modifier discovery in synucleinopathy-relevant circuitry.

## RESULTS

As presented in Figure 1, we outline the end-to-end workflow for DLC model training and subsequent video analysis, from labeled frame selection through model refinement and automated pose extraction (figure generated using BioRender.com).

**Figure 1.**
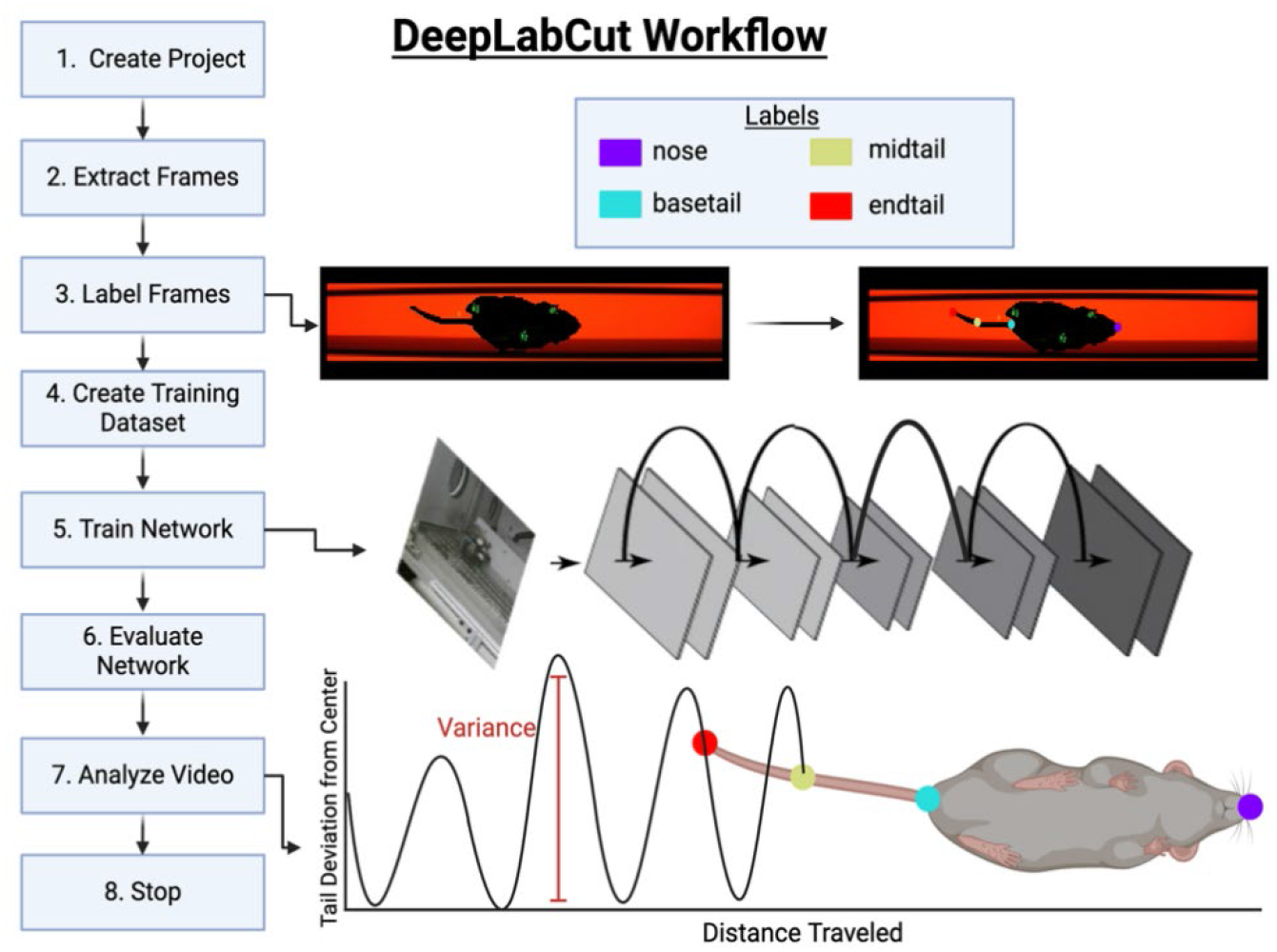
Workflow of DLC model training and video analysis. Figure partially generated using BioRender.com

### Pose labelling of Catwalk video recordings

Two cohorts of male littermate L61-Tg and NTg mice were co-housed in mixed groups up to the age of 12 months, before being assessed by the Catwalk test. After model training (as specified in Methods), we validated body part prediction performance among random frames comparing the position of manually labelled frames against those predicted by DLC. We found that tail-base predictions were significantly more accurate than all other landmarks; therefore, we extracted the tail-base position across all DLC-analyzed videos. Because the Catwalk method orients animals to walk across the training area and in a straight line, the variation in the tail-base position through the animal’s movement can be assessed by the variation in the video y-position, where the Catwalk chamber runs parallel to the x-axis. We observed greater mediolateral (y-axis) variability across the forward direction-of-travel in L61-Tg animals compared to NTg (Figure 2B-C for 12-month-old, Figure 3B-C for 18-month-old). To capture this quality in a single number, we determined the statistical variance in tail-base y-position across all frames within each video. Comparison of this lateral variance found a statistically significant increased degree of lateral variance (quantifying mediolateral instability) in L61-Tg compared to NTg among both 12-month-old (Figure 2A) and 18-month-old (Figure 3A) animals.

**Figure 2.**
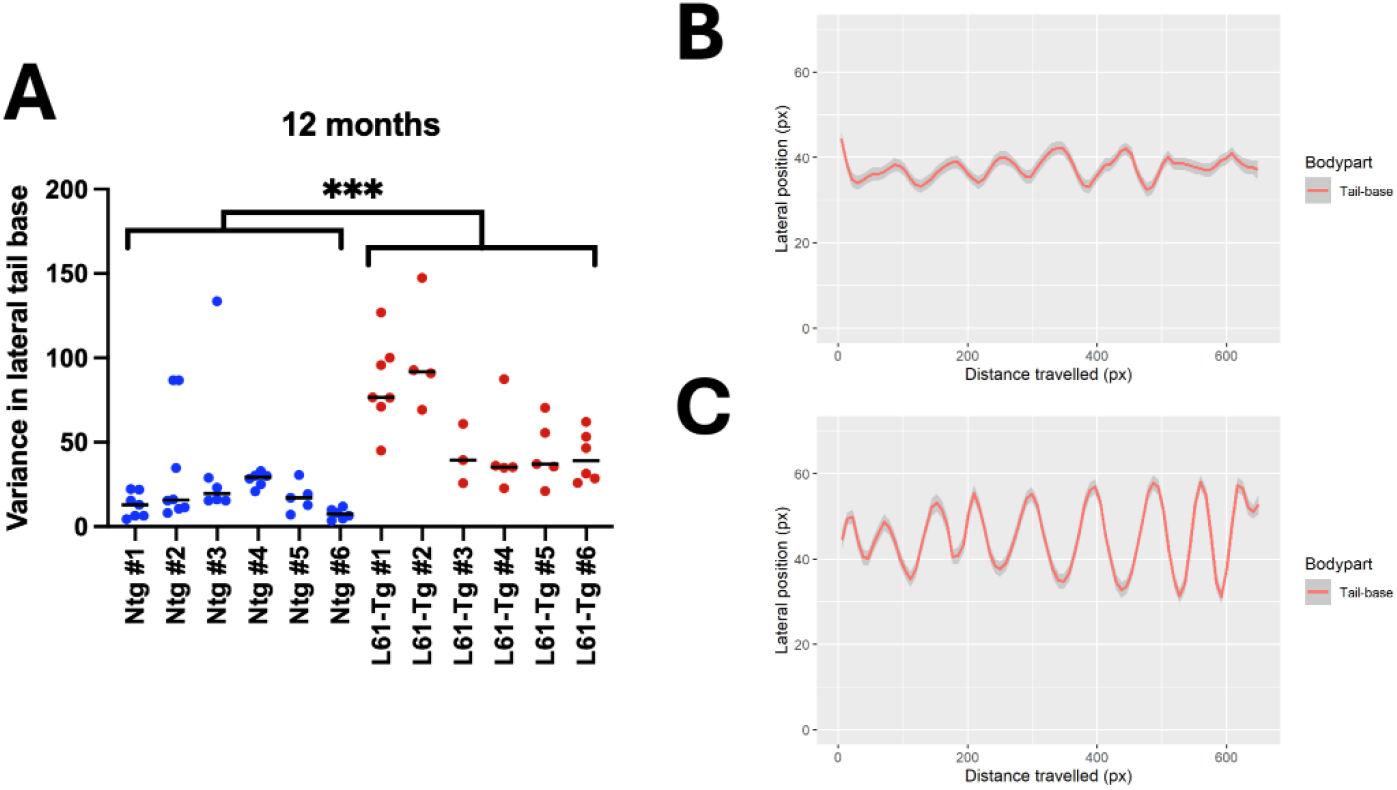
Tail-base lateral movement in 12-month-old L61-Tg vs. NTg mice. (**A**) Dot plot showing lateral variance across NTg and L61-Tg animals; dots represent individual CatWalk runs. Blue dots indicate NTg mice and red dots indicate L61-Tg mice. Bars indicate the per-animal median. Nested two-tailed t-test, L61-Tg vs. NTg: p = 0.0002. (**B–C**) Representative position plots of axial (x, forward direction of travel) versus lateral (y) tail-base position for a single NTg (**B**) and L61-Tg (**C**) DLC-analyzed CatWalk run.

**Figure 3.**
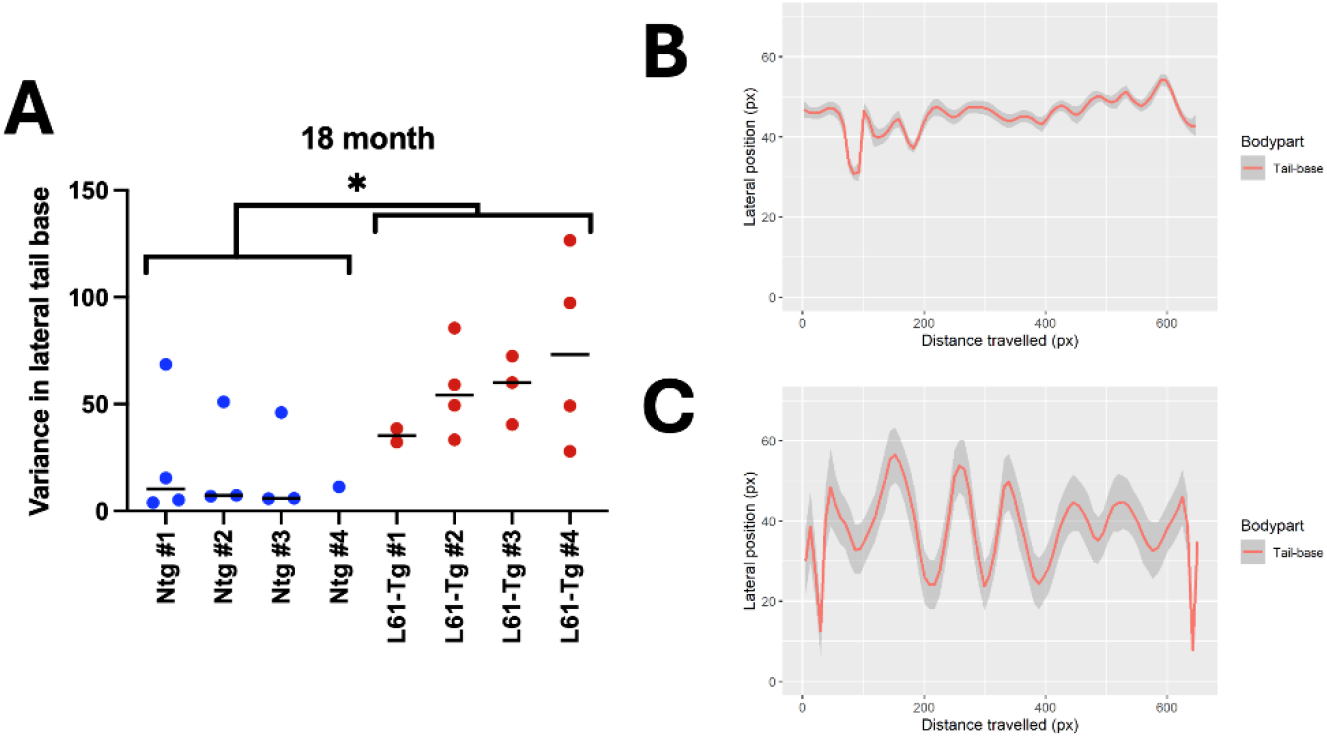
Tail-base lateral movement in 18-month-old L61-Tg vs. NTg mice. (**A**) Dot plot showing lateral variance across NTg and L61-Tg animals; dots represent individual CatWalk runs. Blue dots indicate NTg mice and red dots indicate L61-Tg mice. Bars indicate the per-animal median. Nested two-tailed t-test, L61-Tg vs. NTg: p = 0.0374. (**B–C**) Representative position plots of axial (x, forward direction of travel) versus lateral (y) tail-base position for a single NTg (**B**) and L61-Tg (**C**) DLC-analyzed CatWalk run.

### Motor deficits by Catwalk gait analysis

Analysis using CatWalk followed standard procedures, as specified in Methods. Because the L61-Tg model has bilateral synucleinopathy impairment and is not expected to have asymmetrical motor dysfunction, we combined left and right limb measurements where parameters were recorded unilaterally, such that data represent combined bilateral forelimb (BiF) and combined bilateral hindlimb (BiH). A series of Linear Mixed-Effects models determining prediction by each parameter of L61-Tg vs. NTg status were constructed using a hierarchical structure treating the individual animal as a level within genotype and treating paw side (left vs. right) as a level within individual animal among those unilateral measures, while separate Linear Mixed-Effects models were constructed for non-unilateral measures without paw side as a level. After extraction of p values across all parameter models, we performed Benjamini-Hochberg multiple comparison correction, finding that six parameters (of 52) were statistically significant (p-adj < 0.05) between L61-Tg and NTg animals at 12 months of age. Specifically, L61-Tg animals had reduced front base of support (BOS), increased hind BOS, decreased mean front paw stand index (speed of paw lifting from surface), decreased mean single stance (duration of paw-surface contact), increased cadence, and decreased front paw print width (Figure 4). In contrast, among 18-month-old animals, only hind BOS remained statistically significant after multiple-comparisons correction (Figure 5). Consistent with this, the effect size for hind BOS appeared larger at 18 months than at 12 months (slope 0.589 vs. 0.315).

**Figure 4.**
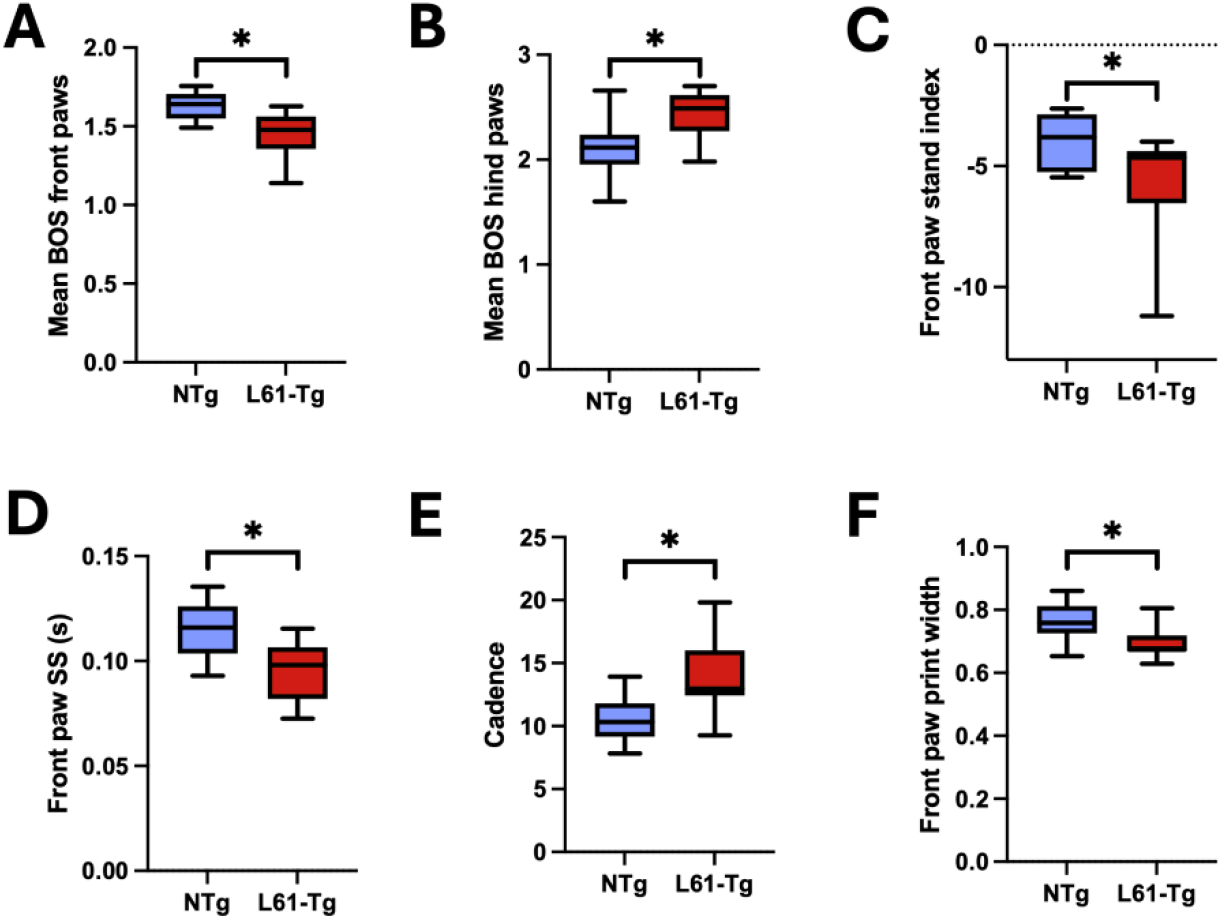
Groupwise boxplots of significantly altered Catwalk parameters between 12-month-old L61-Tg and NTg mice. **(A)** Mean base of support, front paws, **(B)** Mean base of support, hind paws, **(C)** Front paw stand index, **(D)** Front paw single stance (s), **(E)** Cadence, **(F)** Front paw print width. **\***p-adj (Benjamini-Hochberg) <0.05.

**Figure 5.**
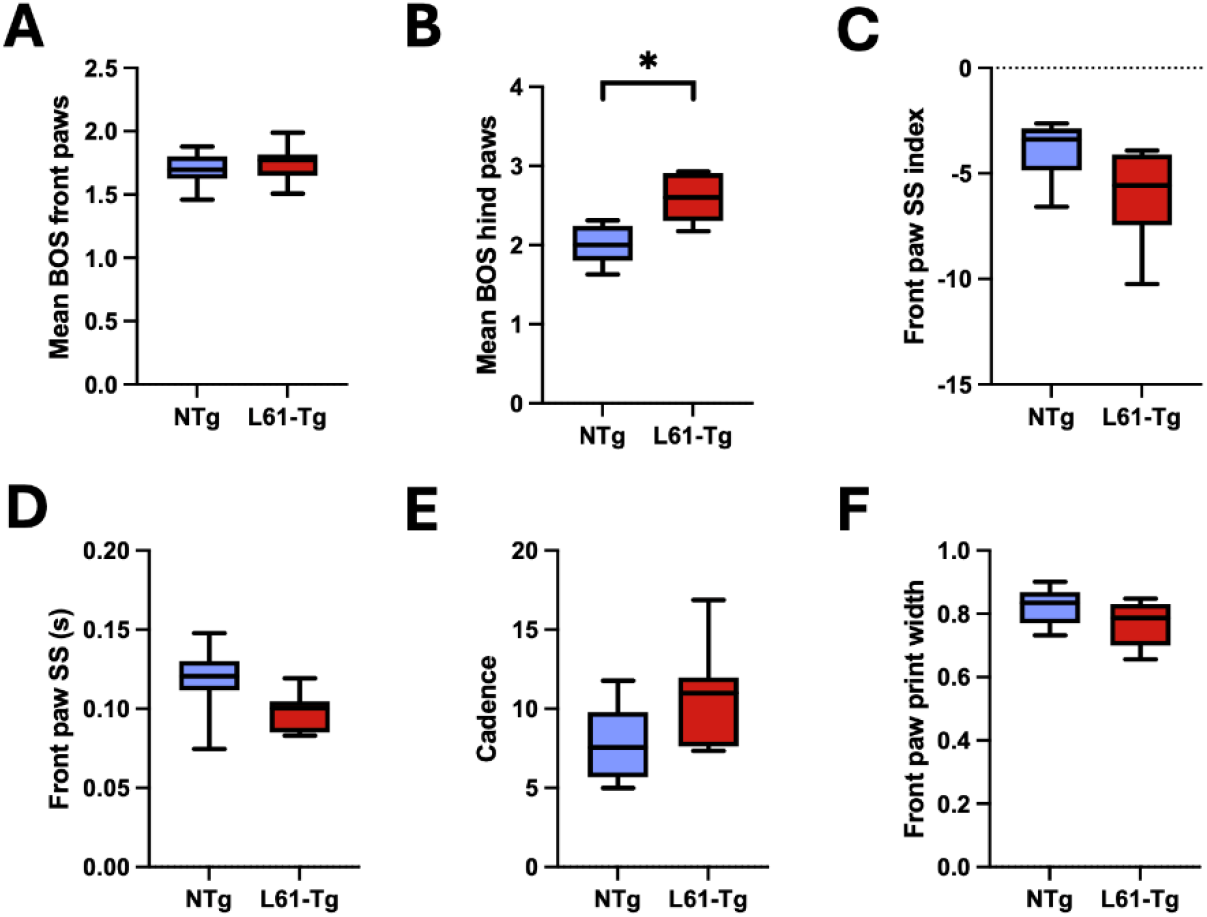
Groupwise boxplots of the six CatWalk parameters identified at 12 months shown in 18-month-old L61-Tg and NTg mice. Only hind base of support differs significantly at 18 months after Benjamini-Hochberg correction. (**A**) Mean base of support, front paws, (**B**) Mean base of support, hind paws, (**C**) Front paw stand index, (**D**) Front paw single stance (s), (**E**) Cadence, (**F**) Front paw print width. *p-adj (Benjamini-Hochberg) <0.05 for hind BOS only.

### Synucleinopathy discrimination of Catwalk parameters and DLC-calculated lateral instability as predictors of motor impairment

To directly compare the performance of the CatWalk parameters to the DLC lateral instability scores (collectively termed “features”), we determined the mean value across runs for each feature before filtering for only mice which were present in both datasets (12-month-old: NTg n=6, L61-Tg n=6; 18-month-old: NTg n=4, L61-Tg n=4). For each feature, we determined receiver operating characteristic (ROC) curves across all animals (12-month-old and 18-month-old) to give a single score measuring feature discrimination which can be interpreted to assess diagnostic discrimination. Such measures which discriminate most strongly between normal and synucleinopathy may therefore represent features which are most sensitive to changes in behavioral phenotype.

These analyses showed, as expected, that several of the predictors which were strongest in the linear model analyses (Figure 4) likewise were strong discriminators between L61-Tg and NTg genotypes (Figure 6). Interestingly, lateral variance and hind-paw BOS were the strongest discriminators by this assessment and each showed complete separation of genotypes in this dataset (ROC-AUC=1.0 for both), in line with the significant genotype differences observed in the prior analyses by nested t tests (for lateral variance) and by linear model (for hind-paw BOS). Given the modest sample size, this “perfect” discrimination should be interpreted as performance within the current dataset rather than a guarantee of out-of-sample generalizability.

**Figure 6.**
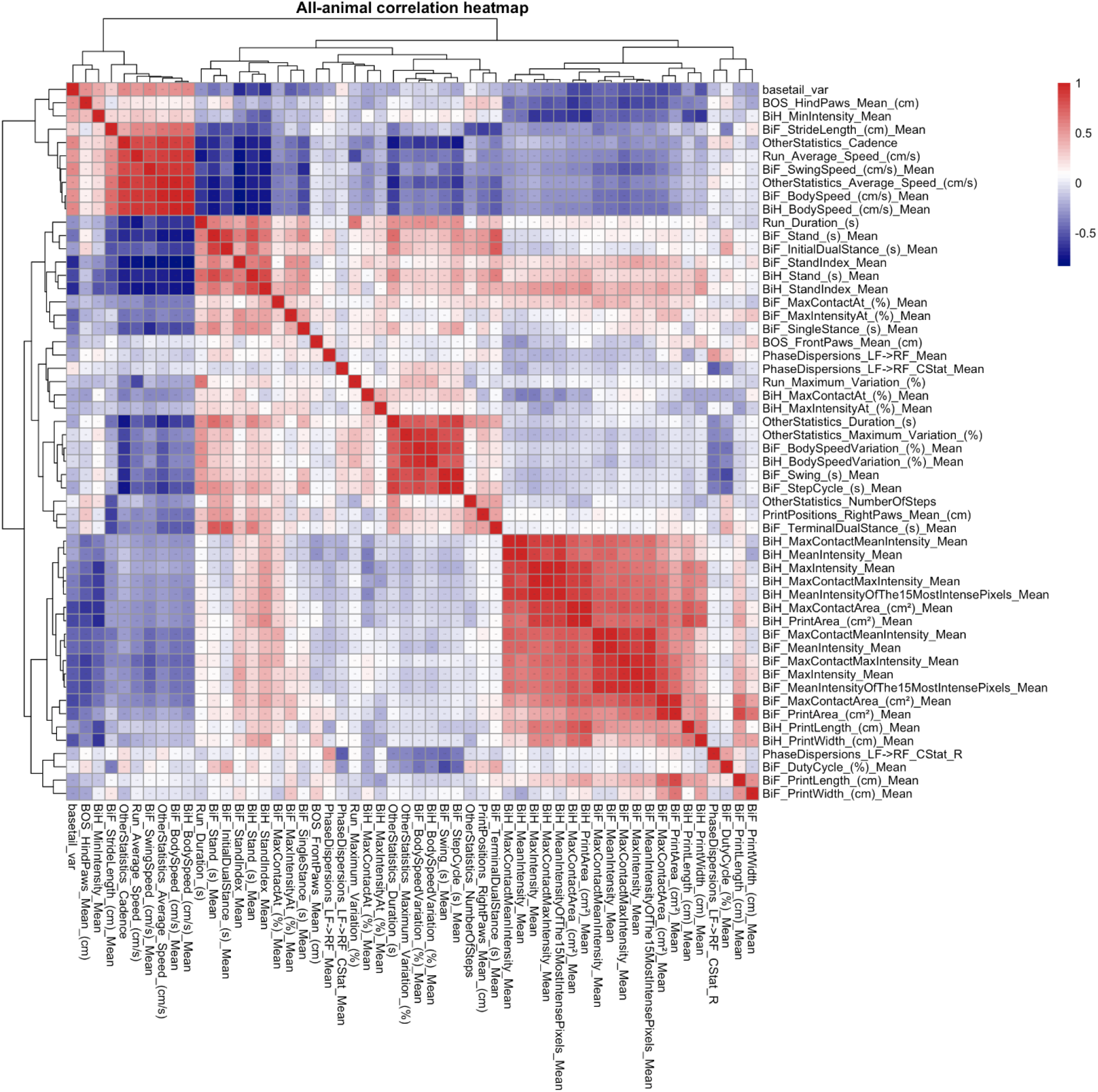
Correlation heatmap across Catwalk parameters and DeepLabCut estimation of lateral variance. Per run (median among bilateral measures), z-scored by standard deviation among all runs and correlated by Pearson method. Parameters with non-normal on invariable distributions discarded. BOS = base of support, BiH = bilateral hind paws, BiF = bilateral front paws.

### Integrative analysis of datasets

Because behavioral parameters are naturally interrelated, we aimed to assess statistically the similarity in differentiation across all parameters. By correlating population-normalized parameter scores across all mice, we identified that parameters occupy distinct clusters (Figure 6), likely indicating that they are measuring either the same gait quality by different means, or are measuring distinct gait qualities which are highly correlated. From this, the two most distinct clusters related to (i) front paw pressure (involving features between “BiF Print Area” and “BiF Max Intensity Mean” in Figure 6), and to (ii) step speed (cluster positively correlated between “BiF Initial Dual Stance” and “Print Positions Right Paws mean”, and then negatively correlated against the cluster between “BiF Stride Length” and “Average Speed”). Interestingly, the two strongest independent predictors, tail base variance and hind BOS, were not within these major clusters, but were correlated closely together (Figure 6, upper left corner). Given the high correlation between these features, we questioned to what extent there might be redundancy (i.e., if tail base variance is determinative of hind BOS, then both hind BOS and tail base variance are two measures of the same gait characteristic (i.e., potentially redundant).

To examine to what extent tail base variance and hind BOS are synergistic rather than redundant, we constructed a random forest (RF) model trained to distinguish L61-Tg from NTg.

Because separation on a per-mouse level could already be achieved with 100% accuracy for both features (ROC-AUC=1 for both, data not shown), we challenged the model to achieve this at a per-run level where there was less distinction between L61-Tg and NTg runs. RF models achieved a per-run accuracy rate of 91.3% (mean of 100 generated models) when all features were provided in the training dataset. Examination of feature importance indicates which features are most frequently used in conjunction with one another and thus contribute most to the overall model accuracy. This showed that tail base variance and hind BOS achieve the highest importance, with almost equal scores (Figure 7). While this indicates that importance is not dominated by either tail base variance or hind BOS, two features which are mutually redundant in prediction may still be assigned similar feature importance if they are very close in their predictive accuracy. To test synergy vs. redundancy conclusively, we then again generated 100 models, but this time removing either tail base variance, or hind BOS, from the feature table to determine whether model accuracy is stable (implying redundancy) or reduced (implying synergy). Such analysis (Figure 8) found that the accuracy (across 100 generated models) decreased from a mean (±95% CI) of 91.27±0.27% when trained with all features, to 88.73±0.36% when trained without tail base variance, and reduced further to 86.79±0.34% when trained without hind BOS, indicating that these measures were synergistic and, within our dataset, that hind BOS on average contributed slightly more to model performance.

**Figure 7.**
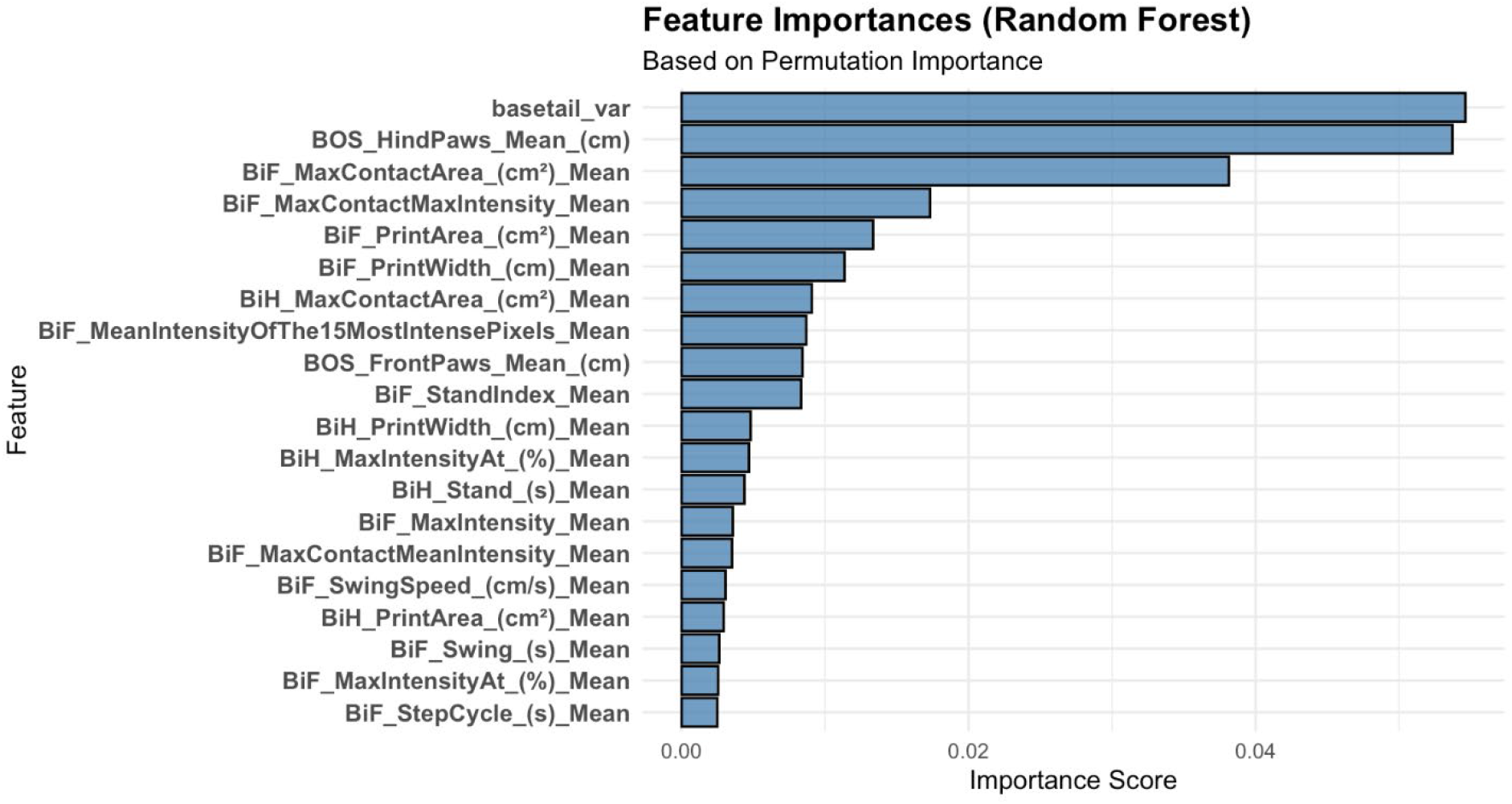
Determination of feature importance within random forest model of the top 20 most important features. Derived from a random forest model trained on all features, with score calculated by permutation importance method. BOS = base of support, BiH = bilateral hind paws, BiF = bilateral front paws.

**Figure 8.**
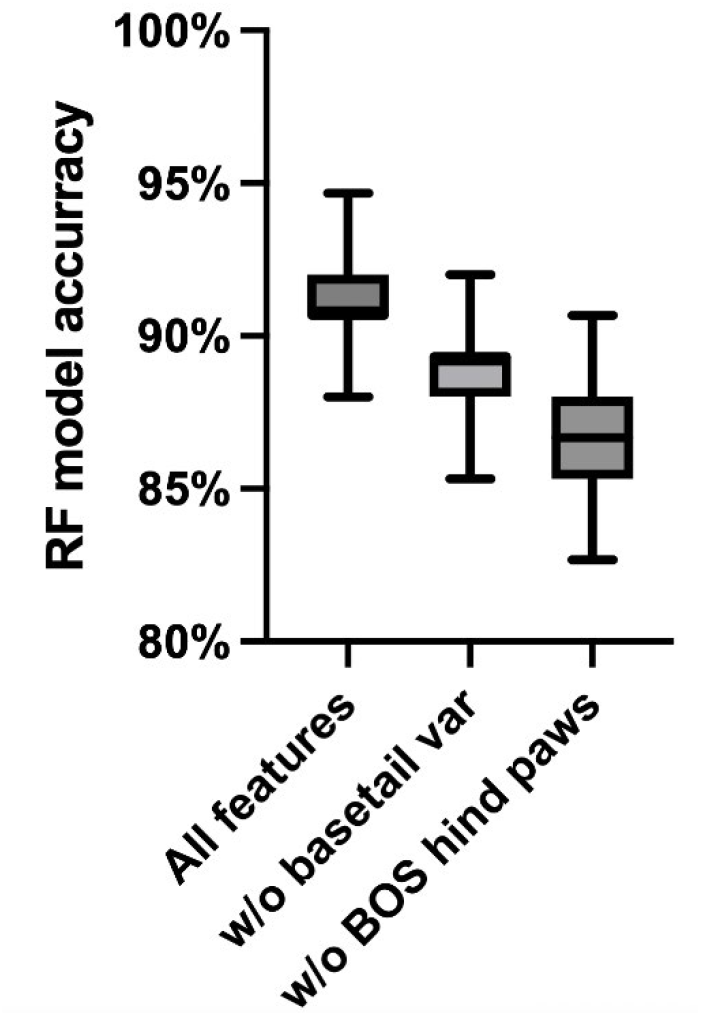
Predictive performance of random forest (RF) models. Models were either trained with data containing all features (“All features”), or all features except for tail base variance (“w/o basetail var”) or all features except for base of support, hind paws (“w/o BOS hind paws”). For each condition, 100 models were generated for each condition to determine a population of model error rates, with each column showing the mean accuracy rate (100% - error rate) with box & whisker indicating minimum, 25^th^ percentile, 75^th^ percentile and maximum values.

## DISCUSSION

This study aimed to develop an objective and scalable method to quantify motor abnormalities in a synucleinopathy-relevant model. The analysis prioritized measures that remain interpretable as gait and postural control. By combining CatWalk XT parameters with markerless pose tracking from the same videos, a compact, stability-focused phenotype was identified that clearly distinguishes L61-Tg mice from NTg littermates. This distinction is particularly relevant for L61-Tg mice, which exhibit a prolonged pre-manifest interval characterized by progressive circuit dysfunction before the onset of dopamine loss and overt disability^15^. Accordingly, the present framework is positioned as a preclinical analogue of “prodromal-to-manifest” monitoring: repeated, quantitative motor sampling that can detect small shifts before conventional disability-like measures change.

A central finding is that a simple DLC-derived metric, tail-base lateral variance during runway traversals, is increased in L61-Tg mice at both 12 and 18 months. In the CatWalk environment, forward progression aligns with the x-axis; thus, increased variance in the y-axis coordinate of the tail base indicates greater mediolateral deviation during locomotion. This measure is directly auditable, as representative coordinate traces reveal the underlying behavior rather than only a summary statistic. The focus on the tail base is technically justified because it was the most accurate landmark in manual-versus-predicted validation, consistent with the principle that DLC output quality is both landmark- and context-dependent and that downstream inference should prioritize reliably tracked points^11^. Notably, because this instability metric is computed across frames within a run, it behaves like a “within-episode” stability readout, which may be especially sensitive to subtle control deficits that are not fully captured by mean gait summaries alone.

CatWalk analysis complements these findings by revealing genotype-dependent gait features at 12 months and a particularly prominent, progressive increase in hind base of support at 18 months. Increased hind base of support is interpreted as a stability strategy and is supported by precedent in parkinsonian mouse phenotyping. For example, in an MPTP murine model^16^, CatWalk gait measures, including base-of-support metrics, track dopaminergic injury severity and correlate with nigrostriatal tyrosine hydroxylase levels, supporting their interpretation as biologically meaningful locomotor readouts rather than arbitrary footprint descriptors^9^. The convergence of two top discriminators from distinct modalities – hind base of support widening and increased mediolateral tail-base variability – supports the conclusion that altered mediolateral stability and compensatory postural strategy are core components of the L61-Tg motor phenotype at these ages. The directionality is also internally coherent: widened hind BOS is consistent with an adaptive “stance-widening” strategy that would be expected when mediolateral control is noisy or unreliable.

The integrated pipeline provides valuable information because CatWalk and DLC assess complementary aspects of locomotion. CatWalk captures contact timing, base of support, and interlimb coordination, while DLC contributes body-level kinematics that help determine whether changes in gait parameters reflect alterations in postural strategy. The lack of clustering of the DLC parameter with Catwalk parameters (Figure 6) indicates it is not simply measuring the same characteristic by other means and even compared to its strongest co-correlating Catwalk parameter, drop-column analysis demonstrates a significant synergistic interaction (Figure 8). Together, these findings support the interpretation that tail-base instability and support-related CatWalk measures, while both being highly sensitive to the synucleinopathy motor phenotype, capture distinct aspects of motor behavior and should be assessed in conjunction rather than competition with one another. Importantly, this “synergy” result is most compelling in the per-run classification challenge, where within-animal variability and run-to-run noise more closely resemble real-world longitudinal monitoring. Thus, the combined feature set does not merely separate groups once; it improves discrimination under a stricter repeated-measures design.

An additional implication of the Results is that the breadth of CatWalk differences appears larger at 12 months than at 18 months (where hind BOS dominates). This pattern should not be interpreted as “recovery” with age; rather, it may reflect (i) reduced power at 18 months after harmonization across modalities, (ii) increased behavioral heterogeneity in older animals, and/or (iii) a shift from multi-parameter gait alteration toward a more specific compensatory stability signature as disease progresses. A related methodological point is that several per-mouse ROC-AUC values reached 1.0; while encouraging, perfect separation in small samples can overestimate generalizability. Future work should thus emphasize pre-registered cross-validation strategies (e.g., leave-one-mouse-out, cohort-to-cohort replication, and threshold-locking) to quantify how well these features transport across batches, experimenters, and timepoints.

Several limitations affect interpretation and suggest directions for future research. First, some CatWalk measures that differ at 12 months, such as cadence and stance-related variables, may be influenced by speed and compliance. This will be addressed by controlling for run speed, for example by using speed as a covariate, speed-matched subsets, or speed-bin stratification. Second, because runway traversals can also be influenced by affective state or exploratory strategy, it will be valuable to test whether the instability signature persists after accounting for anxiety-like behavior or arousal (e.g., by including open field-derived covariates or by quantifying within-run hesitations). Third, the 18-month integrated dataset is small after harmonizing animals present in both CatWalk and DLC (n=4 per genotype), which limits statistical power to detect moderate effects. Thus, it is important to emphasize that the lack of significance at 18 months should not be overinterpreted. Fourth, the current DLC instability readout is two-dimensional and based on a single landmark. Although the tail base tracked most accurately, y-variance could reflect true sway, yaw or heading variability, or intermittent low-confidence frames. Adding a trunk axis landmark set and filtering by DLC likelihood could improve mechanistic interpretation and reduce ambiguity^11^. In addition, explicitly separating “path curvature” from “body sway” (e.g., by quantifying centerline trajectory vs. body-axis angular dynamics) would help distinguish steering variability from axial instability. Finally, linking these gait and pose measures to pathology or circuit readouts in the same animals would strengthen biological relevance, as CatWalk phenotypes have been associated with dopaminergic injury markers in toxin models^9^. Given the prominent mediolateral phenotype here, correlating gait instability with nondopaminergic circuit measures (e.g., cerebellar/brainstem/vestibular-relevant readouts, or spinal/brainstem patterning changes) may be especially informative, consistent with the clinical observation that balance deficits can be only partially dopamine-responsive as discussed below.

This stability-focused phenotype is particularly valuable because postural instability in PD and related conditions is both disabling and clinically significant^17–19^. Falls resulting from balance impairment are common throughout the course of disease and represent a major source of morbidity^17–19^. In population-based cohorts, the postural instability/gait difficulty phenotype independently predicts mortality^20^. Notably, postural control deficits often respond poorly to levodopa, indicating substantial nondopaminergic contributions; quantitative studies have shown that levodopa does not significantly improve balance responses, highlighting treatment-refractory aspects^21^. The physiopathology and functional neuroanatomy of postural instability remain unclear, and recent reviews emphasize that less is known about the underlying neural network dysfunctions than about biomechanics^22,23^. Given these gaps, the present testing strategy may be preclinically important, as it provides objective, scalable, and video-auditable measures of mediolateral instability during walking. This approach enables the dissection of circuit mechanisms and the evaluation of candidate interventions in models such as the L61-Tg mouse model, where progressive synucleinopathy can be studied prior to the onset of late-stage disability. From a translational endpoint perspective, the combination of (i) an interpretable gait strategy (hind BOS widening) and (ii) a direct kinematic instability readout (tail-base lateral variance) provides a compact feature set that is well-suited to therapeutic screening: these measures are automated, repeatable, and can be tracked longitudinally as continuous outcomes rather than threshold-based impairment scores.

Finally, several design expansions would strengthen future studies. (1) Inclusion of females is important for generalizability, given sex-dependent differences in motor aging and synucleinopathy vulnerability. (2) A fully longitudinal design following the same mice across ages would disentangle cohort effects from disease progression and directly estimate within-animal slopes, which are often more sensitive than cross-sectional contrasts. (3) Establishing assay reliability (test-retest stability, inter-operator robustness, and day-to-day variance) would better define the smallest detectable change, a key parameter for intervention studies. Together, these steps would further position the combined CatWalk-DLC pipeline as a rigorous, scalable motor biomarker platform for synucleinopathy-relevant preclinical research.

## METHODS

All animal related procedures and euthanasia were approved by Columbia University’s Institutional Animal Care and Usage Committee. Male mThy1-α-syn mouse line 61 (L61-Tg)^24^ overexpressing wild-type α-syn mice under the expression of the pan-neuronal mThy1 promoter were obtained from The Jackson Laboratories (D2-Tg (Thy1-SNCA)^61Ema/RoriJ^, Jax #038796). Mice were randomly distributed into three cohorts of littermates: cohort one were aged to 12 months and consisted of 5 L61-Tg and 7 Ntg mice male mice in mixed housing, cohort two were aged to 12 months and consisted of 6 L61-Tg and 5 Ntg male mice in mixed housing, and cohort three were aged to 18 months and consisted of 7 L61-Tg and 9 NTg mice. DLC video analysis was conducted on all mice in cohort one, as well as 4 L61-Tg and 4 NTg mice from cohort three, selected at random. All animals were housed in a specific pathogen-free facility with ambient temperature of 18-23 °C and 40-60% humidity and a 12-h light/dark cycle, in a dedicated facility within the Mouse Neurobehavior Core at Columbia University from weening until the end of the study, staying within consistent cages and with cage assignments recorded and randomized across genotypes.

Mice were habituated to the assessment room for 30 minutes before testing. Catwalk apparatus glass walkway and inner walls were cleaned with 70% ethanol and allowed to dry fully before introducing each mouse. Each mouse is trialed until three compliant runs have been recorded, after which it is removed from the apparatus, transferred to a holding cage, the apparatus cleaned thoroughly with 70% ethanol and allowed to dry fully, and the next mouse introduced. Order was randomly determined within cages, which themselves were mixed between genotypes.

Catwalk analytical parameters were determined using Catwalk XT software (version 10.6.608, Noldus 2015). Following automatic exclusion of runs on the basis of a minimum run duration of 0.5s and maximum run duration of 20s, recorded steps were manually filtered for exclusion of non-compliant runs (not crossing, stepping back, exploratory rather than travelling behavior), foot contact periods were determined, with non-compliant periods within runs (e.g. stopping, turning) being excluded but, for those included periods, foot contact being determined by fully automated processing by the Catwalk software. Catwalk parameters were generated and exported at a per-run basis for analysis using R.

A total of 100 Catwalk videos were used for DLC model training, with 68 videos from 12-month-old mice and 32 videos from 18-month-old mice, with videos between 3 and 11 seconds at 100 fps. For both age cohorts, 20 frames were extracted from equally spaced periods across 10 randomly selected videos. Body parts (nose, tail base, tail mid, and tail tip) were labelled manually with the assessor blinded to the mouse identity. From this, DLC was training using default model parameters with 100,000 iterations. Following visual assessment of model performance, labelling was performed onto all frames of all 100 videos to produce tabulated data for estimated position of each body part in each frame. Following import of this data to R, the position of the tail base was extracted and the statistical variance was determined among all frames in each video to compose the ‘tail base variance (var)’ score for the given run.

Catwalk parameters and DLC tail base position parameters were imported for statistical analysis in *R* (v4.3.3), using packages *pROC* (v1.19.0.1), ranger (v0.18.0), faxtoextra(v2.0.0), pheatmap(v1.0.13), lmerTest (v3.2-1), lme4 (v2.0-1), Matrix (v1.6-5), magrittr (v2.0.4), lubridate (v1.9.5), readr (v2.2.0), tidyr (v1.3.2), tibble (v3.3.1), ggplot2 (v4.0.2), tidyverse(v2.0.0) and derivative packages. Statistical tests were performed as described in the results using the relevant functions (if applicable) of the packages specified or (otherwise) using those in base *R* or the standard packages of the distribution (*stats, graphics, grDevics, datasets, utils, methods* and *base*).

